# Co-culture of soil biofilm isolates enables the discovery of novel antibiotics

**DOI:** 10.1101/353755

**Authors:** Chun-Hui Gao, Peng Cai, Zhunjie Li, Yichao Wu, Qiaoyun Huang

**Affiliations:** State Key Laboratory of Agricultural Microbiology, College of Resources and Environment Huazhong Agricultural University, Wuhan 430070, China

**Keywords:** non-ribosomal polypeptide synthase, inter-species interaction, multispecies biofilm, *Pseudomonas*, soil bacteria

## Abstract

Bacterial natural products (NPs) are considered to be a promising source of drug discovery. However, the biosynthesis gene clusters (BGCs) of NP are not often expressed, making it difficult to identify them. Recently, the study of biofilm community showed bacteria may gain competitive advantages by the secretion of antibiotics, implying a possible way to screen antibiotic by evaluating the social behavior of bacteria. In this study, we have described an efficient workflow for novel antibiotic discovery by employing the bacterial social interaction strategy with biofilm cultivation, co-culture, transcriptomic and genomic methods. We showed that a biofilm dominant species, i.e. *Pseudomonas* sp. G7, which was isolated from cultivated soil biofilm community, was highly competitive in four-species biofilm communities, as the synergistic combinations preferred to exclude this strain while the antagonistic combinations did not. Through the analysis of transcriptomic changes in four-species co-culture and the complete genome of *Pseudomonas* sp. G7, we finally discovered two novel non-ribosomal polypeptide synthetic (NRPS) BGCs, whose products were predicted to have seven and six amino acid components, respectively. Furthermore, we provide evidence showing that only when *Pseudomonas* sp. G7 was co-cultivated with at least two or three other bacterial species can these BGC genes be induced, suggesting that the co-culture of the soil biofilm isolates is critical to the discovery of novel antibiotics. As a conclusion, we set a model of applying microbial interaction to the discovery of new antibiotics.

## Importance

Biofilm is an emergent form of bacterial life in natural settings. In a biofilm community, bacteria may employ several different strategies to avoid unwanted neighbors and gain competitive advantages, one of which is the secretion of antibiotics. These antibiotics are bacterial natural products and potential novel antimicrobial drug candidates. In this study, we have described an efficient workflow for novel antibiotic discovery by employing biofilm cultivation, co-culture and multiomics methods. Two novel NRPS-type BGCs were identified in a soil biofilm isolate, namely *Pseudomonas* sp. G7. Furthermore, the metabolite biodiversities of the two BGCs were analyzed by comparing them with those of related *Pseudomonas* spp. Additionally, we provide evidence showing that only when *Pseudomonas* sp. G7 was co-cultivated with at least two or three other bacterial species can these BGC genes be induced, suggesting that the co-culture of several bacterial isolates is critical to the discovery of novel antibiotics.

## Introduction

Biofilms are the most common mode of microbial existence in natural habitats (1). It has been estimated that in natural ecosystems, up to 99% of all bacterial activity is associated with bacteria that are attached to surfaces of and organized into biofilms (2). A biofilm mode-of-life not only gives cells several advantages (3) but also leads to intensive bacterial competition in multispecies biofilm communities (4). To gain an advantage in the competition, a bacterial species may rely on many different approaches, two of which are to inhibit the growth of other species by nutrient competition through the secretion of siderophores and to kill competitors by secreting broad-spectrum antibiotics (5). These approaches have been portrayed as the “pirate” and “killer” approaches, respectively (6). As microbial interactions trigger the production of antibiotics (7), it is logical to conclude that in a mature biofilm community, the dominant strain that is present is more likely to be a producer of antibiotics.

Since pathogen drug resistance is a major threat to human health, finding novel antibiotics is an urgent need. At present, natural products (NPs) are still the main sources of new antibiotics, which are generally secondary metabolites produced from a bacterial biosynthetic gene cluster (BGC) (8). With the revolutionary advances in genomics, metabolomics and other methodologies over last 10-20 years, it has become easy to identify BGCs in a known bacterial genome or meta-genome using bioinformatic tools and resolve the structures of new biomolecules through high-pressure liquid chromatography (HPLC), mass spectrometry and nuclear magnetic resonance (NMR) spectroscopy (9). However, genomic analysis has revealed that many microorganisms have a far greater potential to produce specialized metabolites than is suggested by classical bioactivity screens, as many BGCs in the genome are silent and therefore not expressed under standard laboratory growth conditions (10). This lack of expression makes the screening for novel antibiotic biosynthesis in a bacterial strain a limitation in such a process.

To overcome the limitations in screening and/or try to activate the expression of BGCs, multiple methods have been developed (10). For example, iChip was used to simultaneously isolate and grow soil bacteria under *in situ* conditions (11), which not only greatly increases the throughput of the screening but also enables the acquisition of a large number of bacteria and their products that cannot be cultured using conventional media. In addition, some BGCs are imported into engineered model bacteria and heterologously expressed under strong promoters to promote NP biosynthesis (12). Notably, these efforts are highly dependent on bacterial monocultures while ignoring the essential role of interspecies interactions on inducing BGC gene expression.

Strikingly, methods of co-cultivation, which were initially used to study interspecies interactions in biofilms, are beginning to be applied to the discovery of new antibiotics (7, 13). For example, co-culturing the fungus *Aspergillus nidulans* and 58 actinomycetes species can activate a variety of silent gene clusters in *A. nidulans*, such as polyketide orsellinic acid, lecanoric acid, and the cathepsin K inhibitors F-9775A and F-9775B (14). The coculture of marine invertebrate-associated bacteria has enabled the discovery of a new antibiotic, keyicin (15).

In the current study, we first cultivated soil biofilms using a model system and investigated the microbial succession of biofilm communities during a seven-day cultivation period. From the soil biofilms, twenty-four bacterial strains were isolated. In this case, all of the isolates were potentially dominant species in the biofilm community. Seven isolates belonging to different genera were subsequently picked, and intergenus interactions between the isolates were assessed using a co-culture and the Nunc-TSP Lid system. We found that the presence of *Pseudomonas* isolate G7 had a significant effect in determining whether synergy or antagonism occurred in the four-species biofilm, as the synergistic combinations preferred to exclude this strain while the antagonistic combinations did not. Furthermore, *de novo* (meta-)transcriptomic analysis revealed that novel non-ribosomal polypeptide synthase (NRPS) genes were significantly upregulated in a G7-containing four-species co-culture, which led to the discovery of two novel antibiotic biosynthetic NRPS BGCs.

## Results

### Workflow of this study

We hypothesized that the dominant species in the biofilm communities were more likely to synthesize antibiotics. Therefore, we constructed a model system to produce biofilms of bacteria derived from paddy soil. During cultivation, the community structure of the biofilm was monitored by 16S rRNA gene sequencing. After 7 days of cultivation, the biofilm community was stabilized (see below), and cultivatable bacteria were isolated from the mature biofilm. These isolates are believed be highly competitive in the formation of biofilm, and the isolation of these bacteria allowed us to check the synergistic and antagonistic effects on biofilm formation through the use of co-cultures and high-throughput crystal violet assays (Fig. 1), ultimately identifying the most competitive members.

**Figure 1.**
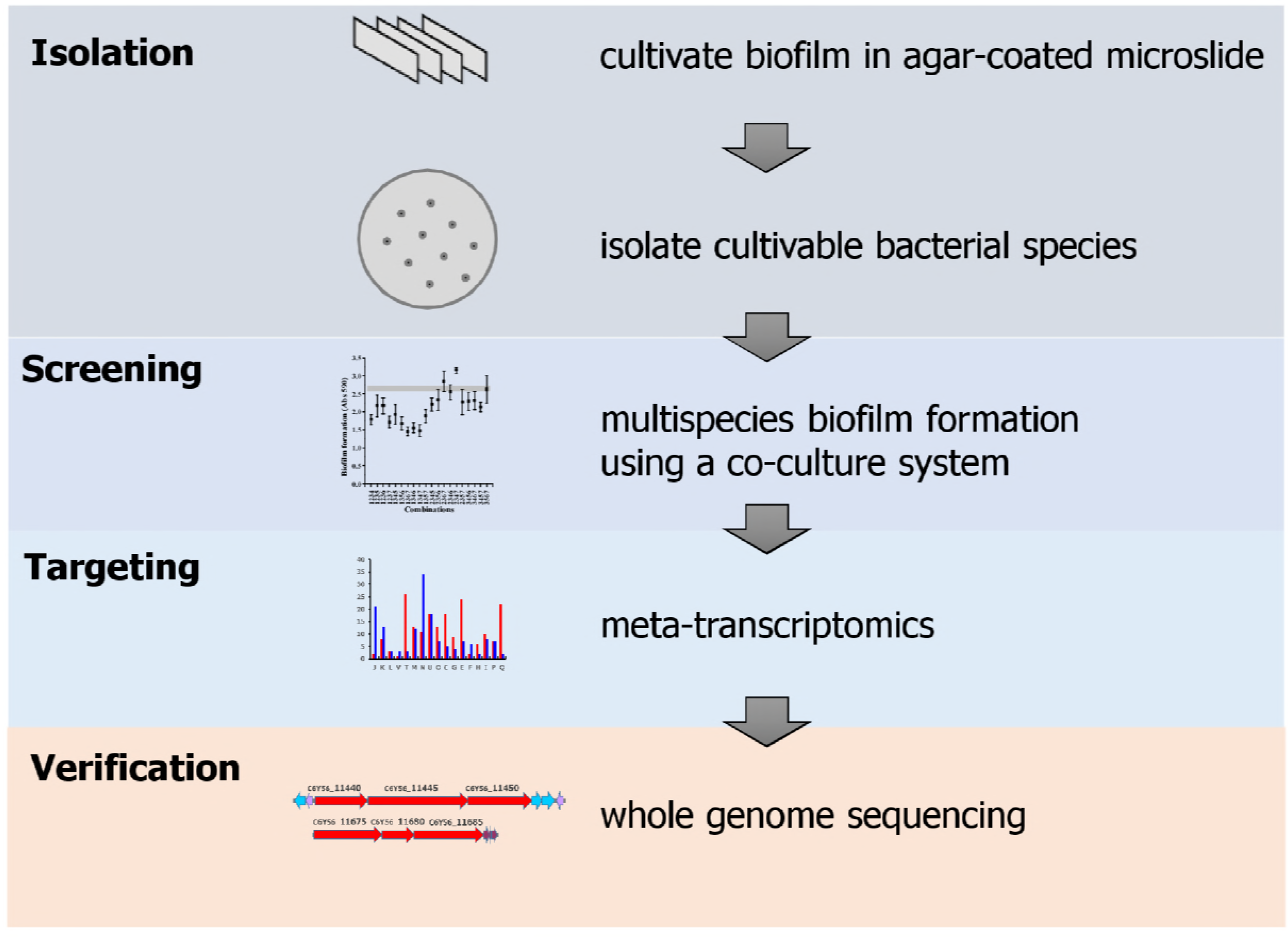
Flow chart of the presented pipeline.

### Bacterial isolates were obtained from biofilm cultivated from a paddy soil microbial community

Biofilm cultivation was initiated from soil microbial communities which had been extracted from the top layer of rice paddy soil, as described in the experimental procedures. During cultivation, the succession of the bacterial community in the biofilm was investigated using 16S rRNA sequencing. The experiment was replicated twice and sampled at 1, 2, 4 and 7 days of cultivation using an autoclaved soil solution as the only source of nutrients. As a result, 269,439 valid sequences of the V4-V5 region of bacterial 16S rRNA genes were produced. In these samples, 256 operational taxonomic units (OTUs), defined by 97% sequence similarity, were classified into 11 phyla, including Firmicutes, Proteobacteria, Bacteroidetes, and Verrucomicrobia (Fig. 2A). The most abundant phylum was Firmicutes and was mainly represented by three genera, namely, *Bacillus, Lactococcus*, and *Enterococcus*. The second most abundant phylum was Proteobacteria and was mainly represented by the genera *Pseudomonas, Massilia*, and *Pseudogulbenkiania*.

**Figure 2.**
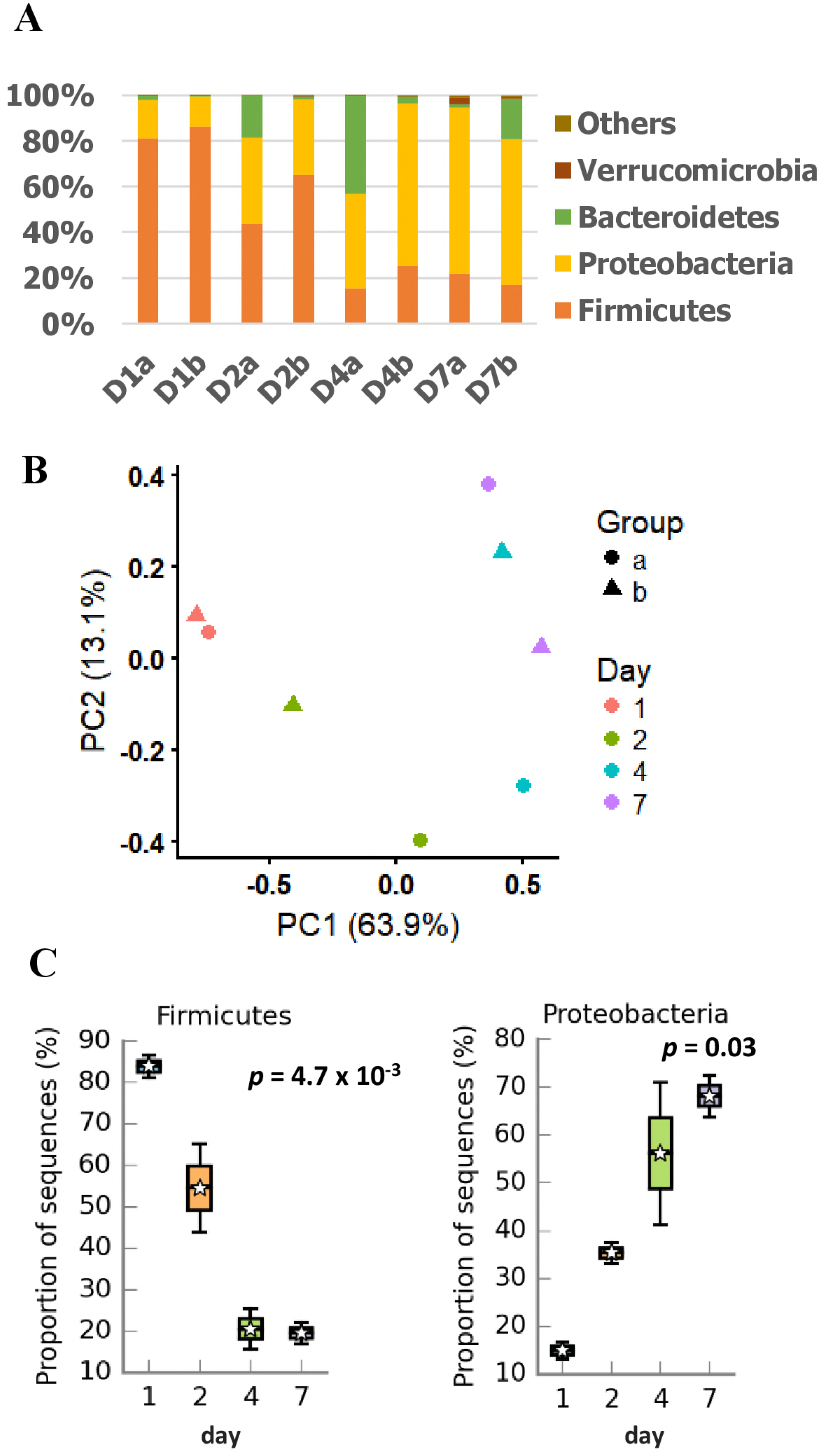
The succession of soil biofilm community. (A) Community structure of biofilm samples over seven days (days 1, 2, 4, and 7). Two replicates were marked by the letter at the end (a, b). Bacterial groups are shown at the phylum level. (B) Principle co-ordinate analysis (PCoA) of bacterial communities. (C) Species that significantly changed in biofilms from days 1 to 7. Significance was showed as p-value on the plots. The mean of the data from each day is shown in the box plot as a star.

Although the culture conditions were very harsh, the alpha-diversity of the biofilm did not decrease but instead slightly increased (Fig. S1). In addition, Principal Coordinate Analysis (PCoA) was conducted to evaluate the similarities between the different samples. The phylogenetic variation, which was measured with UniFrac distances, is displayed in Fig. 2B as a plot of the first two UniFrac principal coordinates. Aside from the two first-day samples, the PCoA did not demonstrate any clustering among the other samples (Fig. 2B); however, time-series samples were separated very well by PC1, which can explain 63.9% of the variation between samples. Notably, the major ordination (PC1) of the day 4 and 7 samples was close, suggesting that the bacteria in the biofilm achieved a relatively stable state at 7 days. From day 1 to day 7, the relative abundance of Firmicutes decreased dramatically, with the proportion changing from ~80 to ~20% of all sequences. In contrast, the abundance of Proteobacteria increased from ~15 to 65% (Fig. 1C), indicating that the dominant Firmicutes species at day 1 were replaced by the Proteobacteria species in biofilms.

Subsequently, twenty-four bacterial strains were isolated from the biofilm communities on day 7 (Table S1). As expected, all of the strains except *Microbacterium* sp. D12 were from dominant phyla in the biofilms, including twelve genera of Proteobacteria and three genera of Bacteroidetes (Fig. 3). These strains were well separated in the phylogenetic tree (Fig. 3), and their coexistence in biofilms suggests that interspecies interactions are ubiquitous among the bacteria.

**Figure 3.**
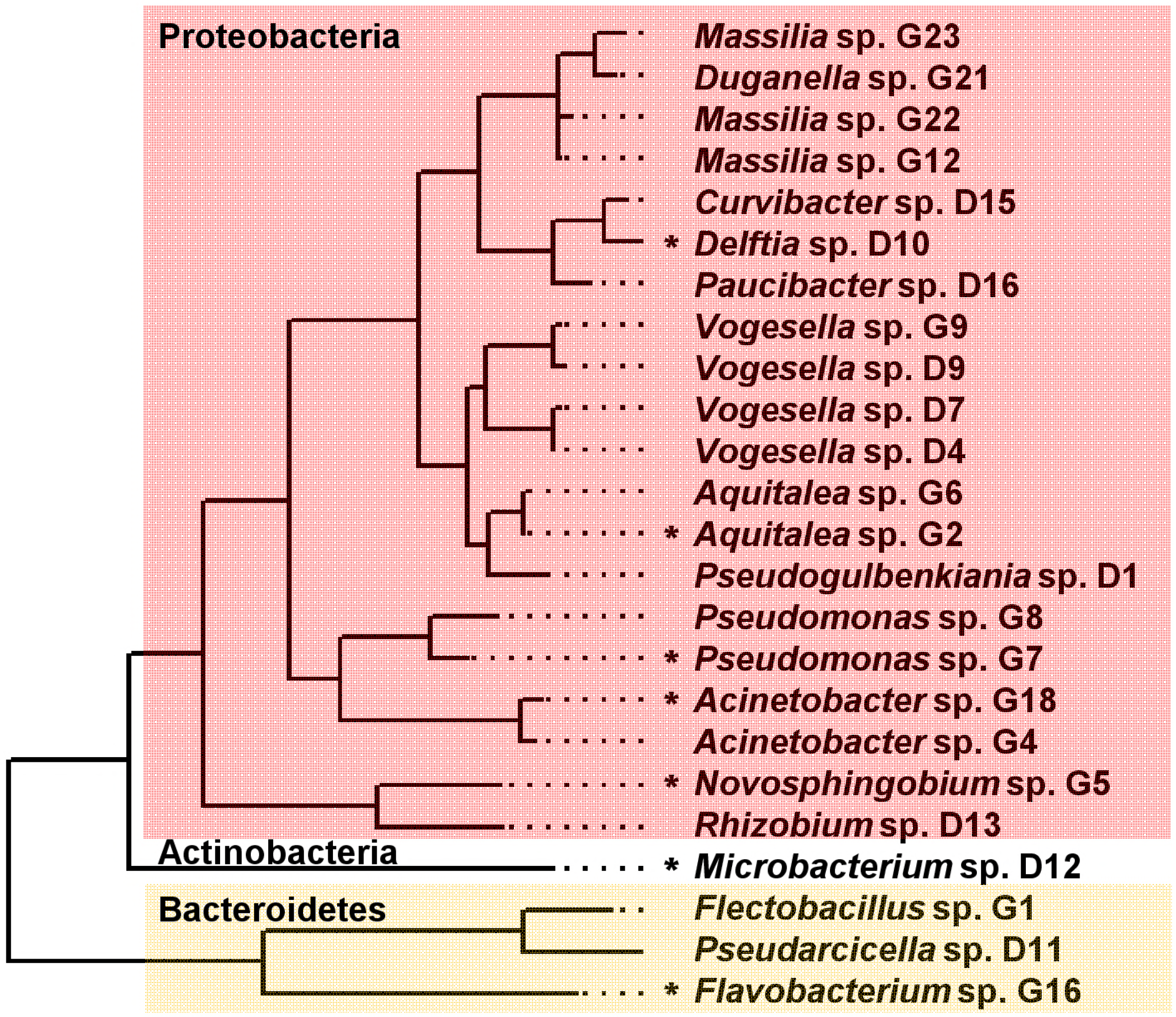
Phylogenetic tree of the 16S rRNA genes of 24 soil biofilm isolates. Phylogenetic analysis was performed by Maximum Likelihood method using MEGA6 (45). The isolates belong to three phylum, showed as different background colours. The asterisk before the strain name indicates that the strain was selected for further co-cultivation and validation. Details of the strain can be found in Table S1.

### Interactions of biofilm isolates revealed the extraordinary role of *Pseudomonas* sp. G7 in multispecies biofilm formation

As stated above, the interactions between different species in biofilm may lead to the activation of antibiotic synthesis, which should be harmful for multispecies biofilm formation. Therefore, it is possible to assess the interactions between different species by quantifying the biofilm biomass. To do this, seven of the isolates from different genera, *Aquitalea* sp. G2, *Novosphingobium* sp. G5, *Pseudomonas* sp. G7, *Flavobacterium* sp. G16, *Acinetobacter* sp. G18, *Delftia* sp. D10, and *Microbacterium* sp. D12 were randomly chosen (Fig. 3) for further analysis. These strains were chosen, in part, because our isolates have relatively rich diversity of species and we expect that more distantly related species may be more likely to compete with each other. These isolates were mixed into four-species co-cultures, and the effects on biofilm formation were analyzed using the crystal violet assay. According to previous data (16), the synergistic or antagonistic effects of co-culture on biofilm formation can be determined by comparing the biofilm biomasses of the co-culture with the monocultures of contained species. That is, when the biofilm biomass of the co-culture is significantly higher than the highest single-species monoculture, the co-culture promotes biofilm formation. When the biofilm biomass of the co-culture is significantly less than the lowest single-species monoculture, the co-culture inhibits biofilm formation.

The biofilm formation of monocultures was initially determined, and the results showed that monocultures biofilm formation varied greatly (Fig. S2), with the biofilm formation of *Pseudomonas* sp. G7 being the highest. Furthermore, a total of 35 different four-species consortia were screened, the biofilm biomasses of twelve combinations were greater than the highest single-species biofilm biomass (Fig. 4). According to the assumptions described above, this result suggested that synergistic effects on biofilm formation exist in the twelve combinations. Of the other 23 combinations, sixteen showed no significant changes while seven showed antagonistic effects (Table S2). Interestingly, most of the synergistic combinations (11 of 12) did not include the strain *Pseudomonas* sp. G7 (Fig. 4). However, most of the antagonistic combinations (6 of 7) did have this strain (Fig. 4, and Table S2). Fisher’s exact test showed that the presence of *Pseudomonas* sp. G7 in co-culture was the most significant factor in determining the fate of biofilm formation (P=0.001), followed by *Acinetobacter* sp. G18 (P=0.006). All the other species were not significant (P>0.05). Consistently, we further found that *Pseudomonas* sp. G7 can form a zone of inhibition on lawns of *Aquitalea* sp. G2 and *Flavobacterium* sp. G16 (Fig. S3B and E), and the *Acinetobacter* sp. G18 can form zones of inhibition on lawns of *Aquitalea* sp. G2 (Fig. S3B), indicating that the two species may be promising antibiotic producers.

**Figure 4.**
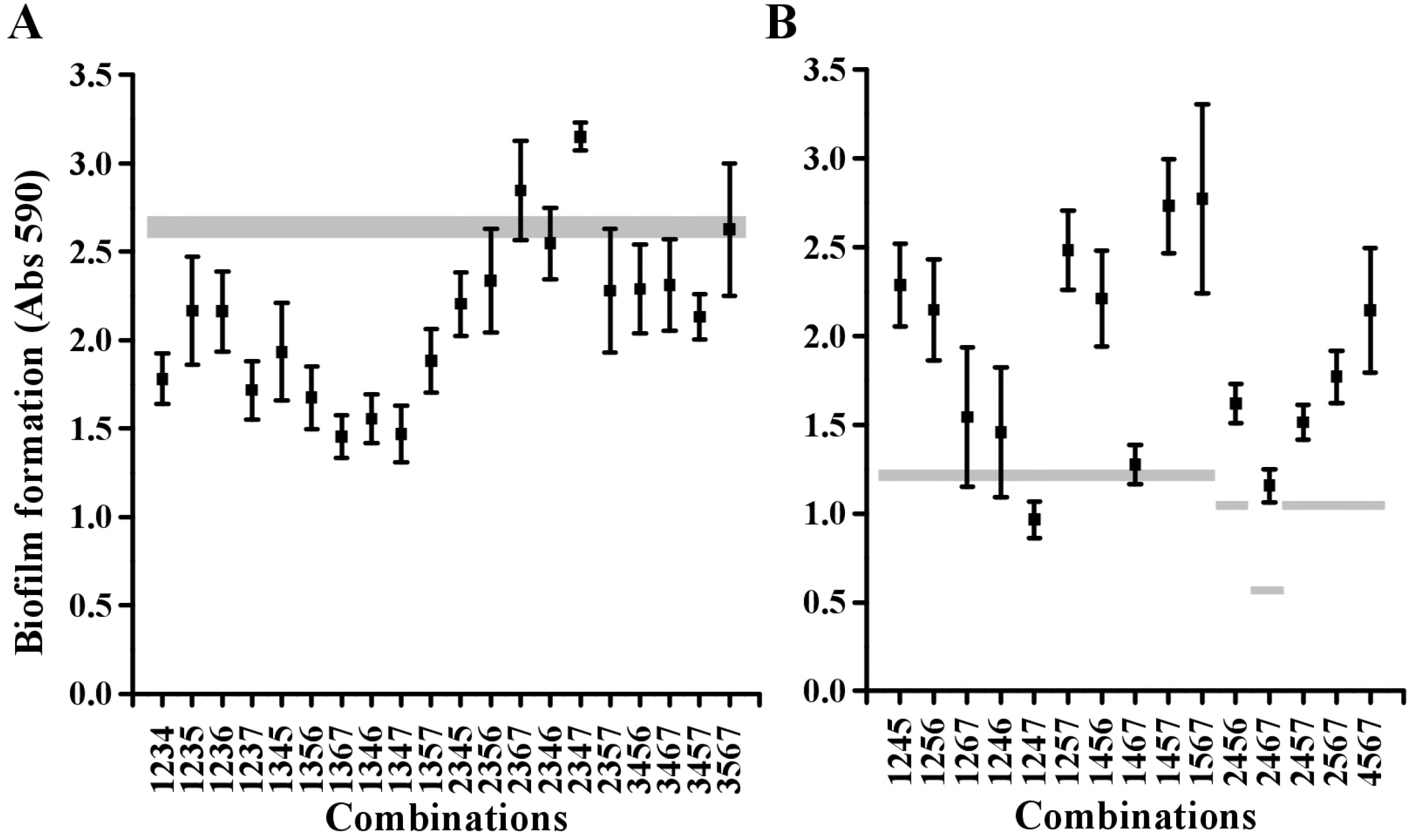
The biofilm formation of four-species co-cultures of seven soil isolates. The observed data points (black dots) were collected by quantifying multispecies biofilm formation of all possible four-species co-cultures using the crystal violet assay. Error bars represent standard deviations of five replicates. The horizontal bars in grey color indicate the amount of monospecies biofilm produced by the best biofilm-forming species that was present in that specific combination. The data obtained from combinations containing strain 3, which is *Pseudomonas* sp. G7, are shown in A, whereas other combinations are shown in B. Strains: 1: *Aquitalea* sp. G2; 2: *Novosphingobium* sp. G5; 3: *Pseudomonas* sp. G7; 4: *Flavobacterium* sp. G16; 5: *Acinetobacter* sp. G18; 6: *Delftia* sp. D10; and 7: *Microbacterium* sp. D12.

### *De novo* meta-transcriptomic analysis revealed the induction of NRPS antibiotic synthase genes in *Pseudomonas* sp. G7 during four-species co-culture

We chose *Pseudomonas* sp. G7 for further confirmation of its capacity to produce antibiotics. To study the molecular mechanisms of *Pseudomonas* sp. G7 in competition with other species, we compared the transcriptomic profiles of a four species co-culture of *Novosphingobium* sp. G5, *Pseudomonas* sp. G7, *Flavobacterium* sp. G16 and *Microbacterium* sp. D12 and monocultures of each individual species using high-throughput mRNA sequencing technology (RNA-seq). Notably, our approach is slightly different from more conventional ones. In the construction of the library, we did not build separate libraries for four mono-cultures but chose to pool them together and construct a single control library (CK). In this way, we only needed to build two transcriptome libraries (without replicates). After sequencing, we first *de novo* assembled the reads, then separated the sequences into the four species using the meta-genomic binning method. Given that we know the species composition in the co-culture system, the use of taxonomic barcodes or phylogenetic marker genes can achieve satisfactory accuracy.

After *de novo* assembly of sequencing reads and meta-genome binning, we finally obtained 4060 unigenes: 1463 unigenes from *Novosphingobium* sp. G5 (3.5 M bps in total), 1280 unigenes from *Pseudomonas* sp. G7 (7.0 M bps in total), 1033 unigenes from *Flavobacterium* sp. G16 (0.14 M in total), and 284 unigenes from *Microbacterium* sp. D12 (0.52 M in total). The variances between the cumulative sizes of the unigenes are attributed to the different abundances of the RNA-seq reads of each species. For example, approximately 95.34% of the total co-culture reads aligned to *Pseudomonas* sp. G7, while only 0.01% of the reads aligned to *Flavobacterium* sp. G16. Due to the lack of RNA-seq reads, the completion of the genome assembly of the other three species was limited. However, this lack of reads from other species also showed that *Pseudomonas* sp. G7 was absolutely dominant in the co-culture.

Most of the unigenes have multiple open reading frames and polycistronic structures. Therefore, genes encoded by these unigenes were predicted prior to the analysis of their expression, as described in the methods section. More than twelve thousand genes were characterized and annotated, and the expression of 461 genes was subsequently found to be significantly changed in the four-species co-cultures compared to their monoculture values (Fig. 5A). The gene expression of *Pseudomonas* sp. G7 was the most affected (421 genes were differentially expressed), followed by those of *Novosphingobium* sp. G5 (37 genes) and *Microbacterium* sp. D12 (3 genes) (Fig. 5). No significant changes in gene expression were found in *Flavobacterium* sp. G16. Of the differentially expressed genes, 383 were successfully annotated by Clusters of Orthologous Groups (COGs), with the number of genes in each functional category being shown in Fig. 5B, grouped by different species.

**Figure 5.**
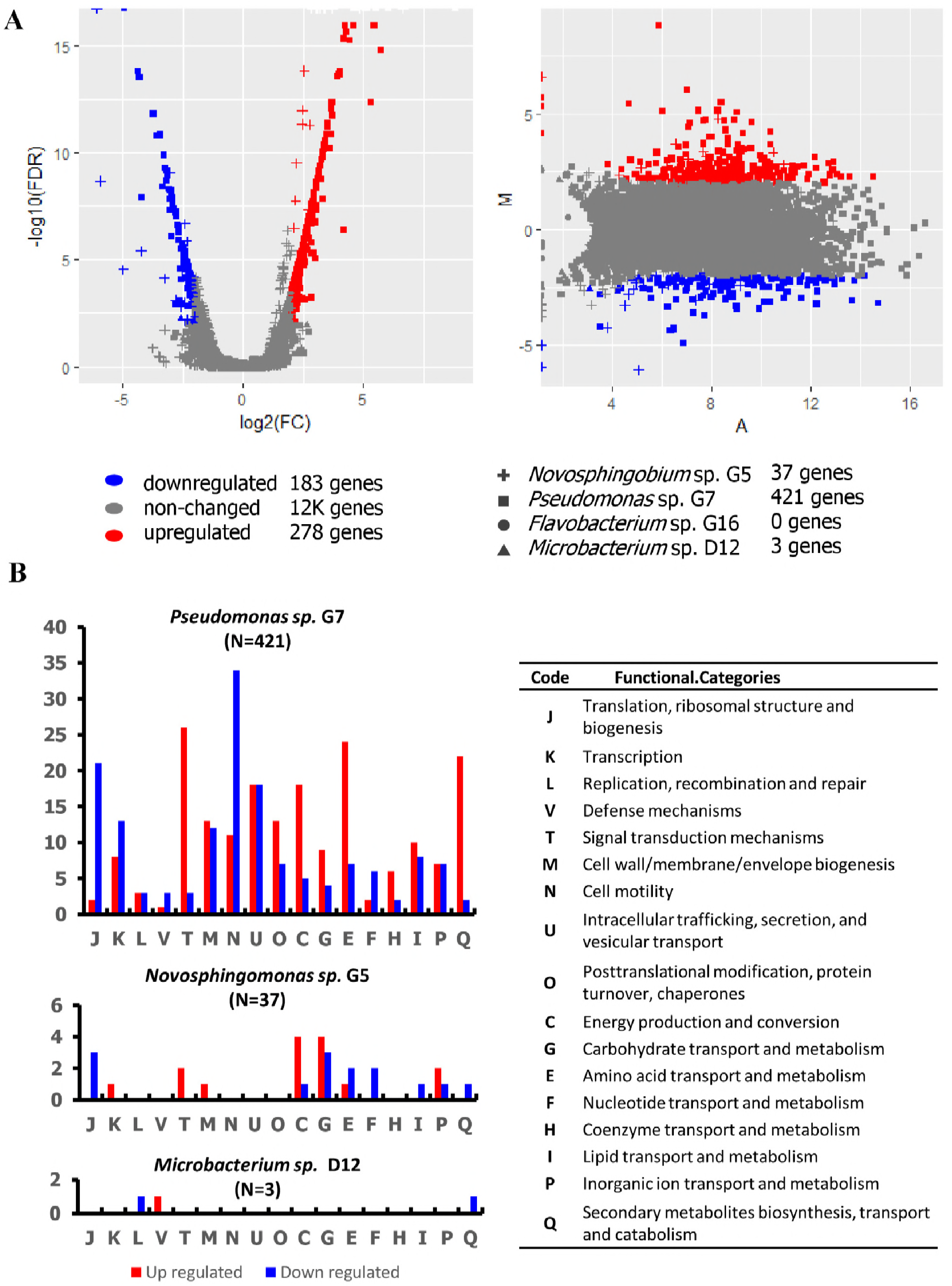
Gene expression changes in four species co-culture. (A) Volcano plot and MA-plot of differentially expressed genes (DEG). Blue points represent down regulated DEG. Grey points represent non-DEGs. Genes that belong to different species were marked with different shapes as indicated by the Figure legend. (B) Functional analysis of DEGs. Number of DEGs in each species are given in the title of each graph. Numbers of COGs are shown in bar plots, and down- (blue) and upregulated (red) genes are represented in different colors. Note that one gene may belong to two or more COG categories. One letter codes are used in the graphs, and the corresponding annotations are shown in the right panel.

As shown in Fig. 5B, most of the genes in the COG category of ribosomal structure (COG: J) were downregulated in both *Pseudomonas* sp. G7 and *Novosphingobium* sp. G5, suggesting that protein translation was inhibited in co-culture. Additionally, multiple metabolic genes (COG: C, G, E, F, H, I, and P) of both species were also significantly changed. In *Pseudomonas* sp. G7, twenty-one transcriptional regulators (COG: K) and twenty-nine signal transduction genes (COG: T) were differentially expressed, implying a complex regulatory mechanism involved in maintaining the activity of the species when in the competition with the other three species. In addition, forty-five genes related to cell motility (COG: N) were differentially expressed in *Pseudomonas* sp. G7, with the majority of these genes, which encode flagellar basal proteins and/or are involved in flagellar biosynthesis pathways, being downregulated. This downregulation suggests that cell motility was inhibited, and biofilm formation was enhanced in co-culture.

Notably, twenty-four genes involved in secondary metabolite biosynthesis, transport and catabolism (COG: Q) were differentially expressed (Fig. 5B), with the majority of non-ribosomal peptide synthetase (NRPS) encoding genes being upregulated more than twenty times in co-culture (Table 1). These NRPS genes were located in five unigenes (Fig. S4). Sequence analysis of these unigenes revealed that they all showed a high sequence similarity (but importantly, were not identical) to a previously identified bananamide biosynthesis gene cluster in *Pseudomonas* sp. DR 5-09 (17). Except for a gap of approximately 1 kb in the *banA* gene, the entire cluster was covered by six unigenes of *Pseudomonas* sp. G7 (Fig. S4). We thus identified that a NRPS antibiotic BGC is present in *Pseudomonas* sp. G7 and highly expressed in co-culture.

**Table 1.** Gene expression changes of NRPS BGCs.

| Meta GeneID | FC | log2FC | P-value | WGS locus_tag | NRPS cluster |
| --- | --- | --- | --- | --- | --- |
| Pseudogene_4428 | 35.6 | 5.2 | 0 | C6Y56_11445 | Paddyleftmide BGC |
| Pseudogene_4429 | 27.0 | 4.8 | 0 | C6Y56_11450 | Paddyleftmide BGC |
| Pseudogene_5485 | 66.5 | 6.1 | 0 | C6Y56_11445 | Paddyleftmide BGC |
| Pseudogene_5486 | 27.0 | 4.8 | 0 | C6Y56_11450 | Paddyleftmide BGC |
| Pseudogene_5487 | 24.7 | 4.6 | 0 | C6Y56_11445 | Paddyleftmide BGC |
| Pseudogene_5488 | 16.1 | 4.0 | 2.16E-14 | C6Y56_11445 | Paddyleftmide BGC |
| Pseudogene_5489 | 27.0 | 4.8 | 0 | C6Y56_11450 | Paddyleftmide BGC |
| Pseudogene_6748 | 27.0 | 4.8 | 0 | C6Y56_11445 | Paddyleftmide BGC |
| Pseudogene_6749 | 33.5 | 5.1 | 0 | C6Y56_11450 | Paddyleftmide BGC |
| Pseudogene_6750 | 26.2 | 4.7 | 0 | C6Y56_11440 | Paddyleftmide BGC |
| Pseudogene_6751 | 35.6 | 5.2 | 0 | C6Y56_11440 | Paddyleftmide BGC |
| Pseudogene_6780 | 24.0 | 4.6 | 1.11E-16 | C6Y56_11440 | Paddyleftmide BGC |
| Pseudogene_303 | 9.1 | 3.2 | 4.09E-10 | C6Y56_11685 | Paddyrightmide BGC |
| Pseudogene_304 | 4.6 | 2.2 | 4.96E-05 | C6Y56_11680 | Paddyrightmide BGC |

Note: Meta GeneID is the coding system that used in Meta-transcriptomic analysis. The
gene ID in the maintext has been replaced by the formal locus\_tag.

### The whole-genome sequencing of *Pseudomonas* sp. G7 revealed two novel NRPS-antibiotic BGCs

Although the meta-transcriptomic analysis showed the presence of a novel NRPS BGC in *Pseudomonas* sp. G7, the sequence of the BGC was incomplete, and therefore, we further performed whole-genome sequencing of this soil biofilm isolate. The complete genome of *Pseudomonas* sp. G7 is 6,336,169 bp, with a GC content of 60.42%, and encodes 5855 genes. The NRPS BGC, which is similar to the bananamide BGC, was then identified by antiSMASH as described in the methods section. Its core region is approximately 35 kb and has three core biosynthetic genes, which are encoded by the ORFs C6Y56_11440, C6Y56_11445, and C6Y56_11450 (Fig. 6A). Notably, an additional NRPS antibiotic BGC (Fig. 6B) was characterized in the area downstream from ORF C6Y56_11450. As annotated by antiSMASH, the two NRPS BGC are both synthetic antibiotics that have not been reported before. Thus, we named their products paddyleftmide and paddyrightmide, respectively.

**Figure 6.**
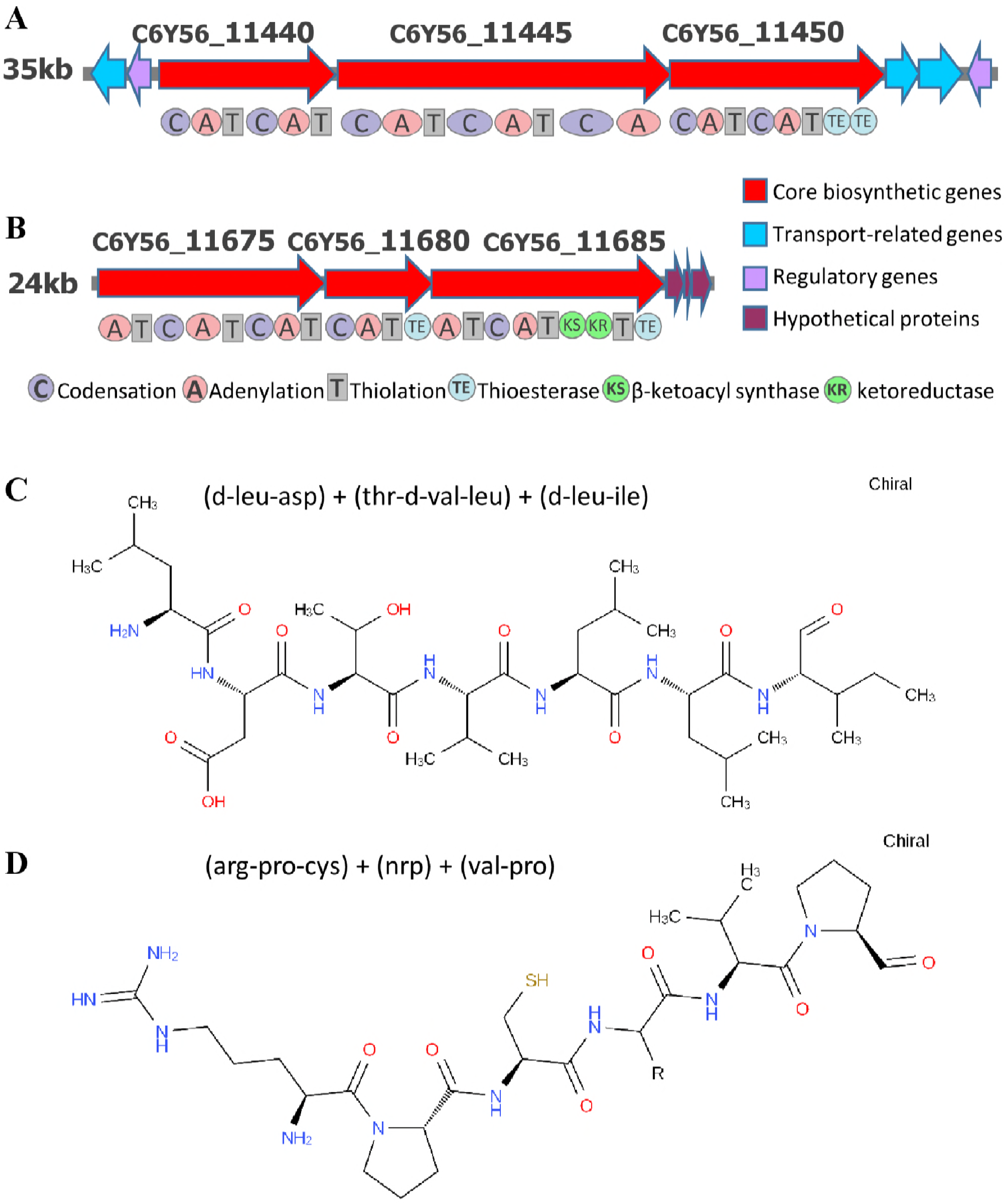
Novel antibiotic biosynthesis gene cluster and predicted molecular structure in *Pseudomonas* sp. G7. (A-B) Genomic structure of thee BGCs. Different kinds of genes were annotated with different colours and function domains in NRPS genes were showed beneath gene symbols. (C-D) Predicted structure of BGC products. The most likely peptide chain composition of the compound is shown on above.

Paddyleftmide was predicted to consist of seven amino acid residues of (d-leu-asp) + (thr-d-val-leu) + (d-leu-ile) (Fig. 6C). The first two amino acids were added by C6Y56_11440, the second three amino acids are added by C6Y56_11445, and the last two amino acids are added by C6Y56_11450 (Fig. 6A). Likewise, paddyrightmide was predicted to have six amino acid residues of (arg-pro-cys) + (nrp) + (val-pro) (Fig. 6D). The first three amino acids were added by C6Y56_11675, the second amino acid were added by C6Y56_11680, and the last two amino acids were added by C6Y56_11685. Notably, both paddyleftmide and paddyrightmide are chiral compounds, and their real structures may be quite different from this *in silico* prediction.

Although we found that the two newly discovered antibiotic BGCs were not included or reported in the MIBiG database (18), they still had some similarities with several publicly available genomes at the sequence level. As shown in Table 2, all of the related genomes were *Pseudomonas* species. Among the genomes was the *Pseudomonas fluorescens* Pf0-1 strain, which is a model microorganism for *P. fluorescens* studies (19). In addition, *P. fluorescens* strain BW11P2 was a relatively early isolate *of Pseudomonas* (20); however, its ability to synthesize bananamide was only recently identified (17). In addition, several other strains were isolated in recent years. All of these bacteria were described as being isolated from soil samples, except for *Pseudomonas moraviensis* strain BS3668, which did not have information available.

**Table 2.** The genome and general information of related *Pseudomonas spp*.

| Strain | Accession | Location | Date | Reference |
| --- | --- | --- | --- | --- |
| *Pseudomonas* sp.<br>DR 5-09 | CP011566 | Daecheong,<br>South Korea.<br>Plant root. | Sep,<br>2012. | Han,J.-H. and Kim,S.B.<br>Unpublished. |
| *Pseudomonas* sp.<br>MS586 | CP014205 | Cotton field,<br>USA | Aug,<br>2012 | Lu,S. and Deng,P.<br>Unpublished. |
| *Pseudomonas*<br>*moraviensis* strain<br>BS3668 | LT629788 | NA | Feb,<br>2016 | Varghese,N. and<br>Submissions,S.<br>Unpublished. |
| *Pseudomonas*<br>*fluorescens* Pf0-1 | CP000094 | Agricultural<br>loam soil | 1988 | (19) |
| *Pseudomonas*<br>*fluorescens* strain<br>BW11P2 | KX437753 | Banana<br>rhizoplane, Sri<br>Lanka | 1992 | (17, 20) |
| *Pseudomonas* sp.<br>G7 | CP027561 | Paddy soil,<br>China | 2016 | This study. |

We compared the phylogenetic similarities of paddyleftmide and paddyrightmide BGCs with similar BGCs in other *Pseudomonas* species. In this analysis, the amino acids of the core biosynthetic genes were used in a multilocus sequence analysis as described in the methods section. The paddyleftmide BGC was phylogenetically different from all other analogs and was not assigned to an end clade of the evolutionary tree (Fig. 7A). However, the distance matrix clearly showed that the differences between *Pseudomonas* sp. G7, P. *fluorescens* Pf0-1 and *P. moraviensis* strain BS3668 is much greater than the differences between *Pseudomonas* sp. G7, *P. fluorescens* strain BW11P2, *Pseudomonas* sp. MS586 and *Pseudomonas* sp. DR 5-09 (Fig. 7B). Likewise, the paddyrightmide BGC is quite unique as well (Fig. 7C). The distance between the BGC and all other BGC analogs is greater than the distance between any of the other analogs (Fig. 7D). However, the absolute difference is more significant for the paddyleftmide BGC, as can be seen when the distance matrixes were colored with an identical color palette (Fig. 7B and 7D). As a conclusion, the two BGCs found in *Pseudomonas* sp. G7 are novel antibiotic producing elements, reflecting the biological diversity of the secondary metabolites of *Pseudomonas spp*.

**Figure 7.**
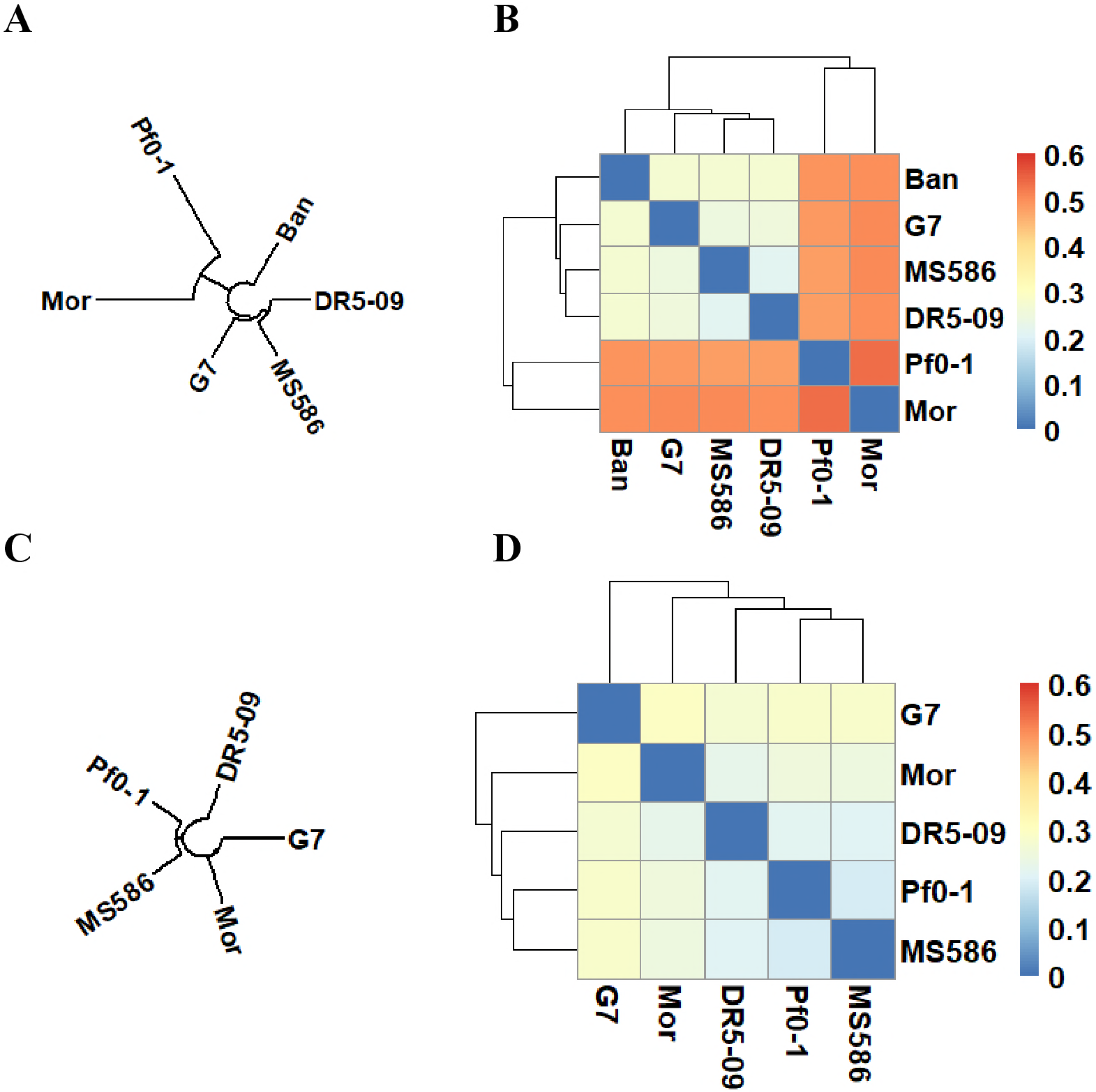
Comparative analysis of the paddyleftmide and paddyrightmide BGC with other BGCs found in related *Pseudomonas* spp. Phylogenetic analysis of paddyleftmide (A, B) or paddyrightmide (C, D) BGC with five or four similar BGCs from related *Pseudomonas* spp., which have been listed in Table 2. Maximum likelihood phylogenetic tree (A, C) was inferred by multilocus sequence analysis of core synthetic genes and the pairwise phylogenetic distance between them (B, D) were shown in heat map with same color theme.

### Reverse verification of the effects of co-culture promoting antibiotic biosynthesis using qRT-PCR

After obtaining the whole-genome sequence of *Pseudomonas* sp. G7, it is easy to find that the upregulated NRPS genes in the Meta-transcriptomic analysis are all from paddyleftmide and paddyrightmide core biosynthetic genes (Table 1). Although we came to the above conclusions through meta-transcriptomic analysis and further whole-genome sequencing, it is still doubtful whether this process is a coincidence; that is, if we choose different combinations of co-cultures, can we still get the same results?

To answer this question, we used qRT-PCR to detect multiple co-culture systems containing *Pseudomonas* sp. G7. The results of the experiments finally demonstrated that even if different co-culture combinations were chosen, we could still find that the NRPS gene was induced by the species-species interactions. Fig. 8A shows the gene expression of the paddyleftmide synthetic gene (C6Y56_11440) in the *Pseudomonas* sp. G7 mono-culture, four two-species co-cultures, six three-species co-cultures, and four four-species co-cultures (all the co-cultures contain *Pseudomonas* sp. G7). Compared with monoculture, the expression of the paddyleftmide synthetic gene was downregulated in all the two-species co-cultures but was mostly upregulated in the three-species co-cultures and was upregulated in all the four-species co-cultures. In general, it can be found that the co-cultures containing at least two species effectively induce paddyleftmide biosynthetic gene expression, and the more species the co-culture contains, the more obvious the upregulation is. Similar conclusions were obtained when measuring the gene expression of the paddyrightmide biosynthetic gene (C6Y56_11675, Fig. 8B). Although there are differences in the expression of the antibiotic synthetic genes in different four species of co-cultures, paddyleftmide and paddyrightmide will eventually be identified in *Pseudomonas* sp. G7, regardless of the combination of species we chose in the four species of co-culture. This result reversely validates that our workflow is robust.

**Figure 8.**
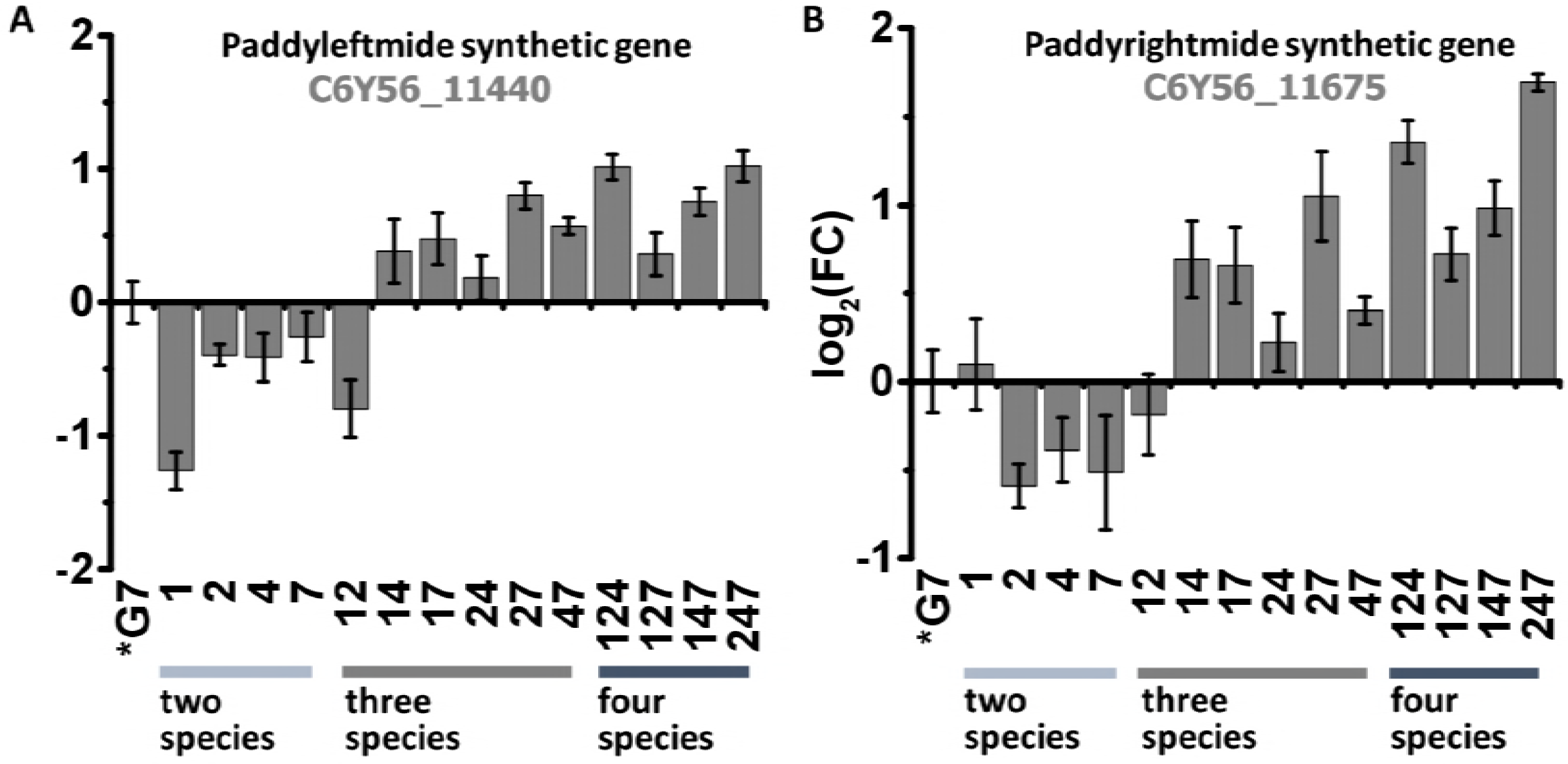
The induction of NRPS synthetic gene in co-culture revealed by qRT-PCR assays. The induction of paddyleftmide (A) and paddyrightmide (B) synthetic gene in *Pseudomonas* sp. G7 monoculture, two-, three- and four-species co-culture were showed. Gene expression was normalized by 16S rRNA gene of *Pseudomonas* sp. G7, and changes were calculated by the 2^−ΔΔCt^ method. Errorbars represent standard deviations of three replicates. The species contained in the co-cultivation show as numbers in horizontal axis, and the numbers corresponding to the species are the same as in Fig. 4.

## Discussion

This study describes a workflow for screening novel antibiotics from soil using biofilm cultivation, mixed co-cultures, and meta-omics techniques. Following this workflow, we demonstrate its effectiveness with the discovery of two novel antibiotic BGCs, which are named paddyleftmide and paddyrightmide, from a paddy soil community.

We used flow cells to culture the biofilm of soil microbial communities. The species present in these biofilms were all derived from soil microbial communities, but due to the heterogeneity of the soil, these microorganisms may not have reflected in the natural environment. Despite the high biodiversity and biomass of living bacteria in soils, only approximately 10^−6^% of the soil surface is covered by microorganisms (21). Therefore, soil bacteria can only contact their close neighbors. The estimated interaction distances of most soil microbes occur over distances no greater than a few tens of micrometers (22). To allow these bacteria to contact and compete with each other, it is necessary to artificially mix them and grow them together. This can be achieved through the application of *in vitro* model systems where an environmental sample is used as inoculum, or to isolate biofilm-forming strains from specific environments for use in co-cultures (23, 24).

By mixing the bacterial species together and fostering competition between species, highly competitive species can be isolated, which will more likely be producers of antibiotics. The 256 OTUs identified in the biofilm samples are at a comparable level to those of a microscale single bacterial community in bulk soil, which is, on average, limited to fewer than 100 species and tens of hundreds of microbial cells (25). However, this number of OTUs is much smaller than that of soil microbiomes, whose total OTUs may range from 3,731 to 27,147 (26–28). With such a biofilm cultivation process, highly competitive species are enriched for in the biofilm community, and therefore, the efficiency of subsequent screens can be improved.

Soil is believed to be the most complex biomaterial (21) and have the most vast microbial biomass and biological diversity (29) on the planet. About two-thirds of all naturally derived antibiotics currently in clinical use, as well as many anticancer, anthelmintic, and antifungal compounds, are produced mostly by the soil-dwelling Actinobacteria, in particular by species of Streptomycetes (30). However, in our biofilm community, the Actinobacteria were not the dominant species. This may be due to culture conditions, since we used autoclaved soil water as the only source of nutrients and cultivated biofilms in an open flow model system. The resulting conditions for biofilm cultivation are aquatic, aerobic, and have a low supply of nutrients, which are not conditions suitable for the growth of Actinobacteria. However, the biofilm community structure at day 7 (Fig. 2) is similar to that of a flooded paddy soil (31) or wetland (32), in which the microbiomes are dominated by Proteobacteria as well. Changing the cultivation conditions may result in the formation of different biofilm communities and thus enable the discovery of potential antibiotics from other species.

Recent studies have shown that microorganisms in the soil are different from those in the air and industrial environments, and they mainly form biofilms in a cooperative manner (16, 33). However, in this study we showed that if the soil microorganism can produce antibiotics, it will significantly affect the formation of biofilm in multispecies co-cultures. Of the 20 combinations containing *Pseudomonas* sp. G7, only one biofilm biomass increased when compared to the monocultures, and of the 7 microbial combinations that lowered biofilm biomass, 6 contained the isolate (Fig. 4, and Table S2). Most of the synergistic combinations (11 out of 12) did not include the *Pseudomonas* sp. G7 strain, and most of the antagonistic combinations (6 out of 7) had this strain (Fig. 4, and Table S2). The presence or absence of *Pseudomonas* sp. G7 significantly influenced the biomass of co-cultured multispecies biofilm (P = 0.001). At the same time, the presence of *Acinetobacter* sp. G18 also has a significant effect on the formation of multispecies biofilm (P = 0.006). Consistently, both of these bacteria form zones of inhibition when grown on the lawns of other species (Fig. S3). The *Pseudomonas* sp. G7 antibiotic synthesis gene cluster was been identified (Fig. 6) and it is likely that *Acinetobacter* sp. G18 also contains unknown antibiotic BGC. This evidence showed that it is feasible to screen for antibiotic-producing bacteria by comparing changes in biomass of multispecies co-cultured biofilm.

We believe that the key to the discovery of new antibiotics is to stimulate their biological activity. If these antibiotics are only encoded by microbial genome but not expressed, no antibiotic will be produced, and the difficulty in screening for these compounds will be enormous. This article presents a method to elicit the biological activity of antibiotics that are capable of being synthesized by microorganisms as well as a method of co-cultivation to identify these antibiotic-producing bacteria. Notably, based on our results, the co-culture may need at least three species to achieve sufficient stimulation to elicit antibiotic gene expression (Fig. 8). Co-cultivation and the measurement of biofilm biomass with crystal violet staining are low-cost, scalable methods (Fig. 1). High-throughput screening can easily be achieved using instruments such as multiwell microplates, loading robots, and microplate readers (34, 35).

When using conventional methods to study gene expression changes in co-culture systems, a reference genome of each species present in the co-culture is required and many mono-culture transcriptomic libraries are constructed as controls (36). By comparing the transcriptomes of mono-cultures and co-cultures, differentially expressed genes can be identified. In this context, identifying gene expression changes in a four-species co-culture may require the genomic sequencing of four species, four separate mono-culture transcriptome sequences, and one transcriptome sequence of the co-culture. After sequencing, the bioinformatics pipeline for transcriptomic analysis with reference will be utilized and the expression changes in the co-culture system can be obtained. This process is, obviously, very costly. Some other methods, such as designing species-specific microarray chips (37), or separating the species present in co-culture by molecular markers (38), also have the same shortage and are not applicable for novel bacterial isolates without a preexisting knowledge of their genome. In this study, we used an alternative method of meta-transcriptome sequencing by combining the four mono-cultures together to construct a single transcriptome library (control). The control library was then sequenced in parallel with the transcriptome library of the four-species co-culture, followed by *de novo* assembly of meta-transcripts. By genome binning, the assembled contigs can be assigned to 4 different species. Then, the gene expression differences in each species can be determined. In this framework, it is only necessary to construct two transcriptome libraries, one for mixed mono-cultures of the controls and the other for the co-culture, decreasing the overall sequencing cost by more than 4 times. Notably, we can still identify the upregulation of antibiotic BGC genes and determine the species in whose genome the BGC is located. These results showed that this improvement of cost does not substantially effect the efficiency of the screening and is therefore beneficial for large-scale screening.

It is noteworthy that this study adopted the process of first separating the cultivatable species and then co-cultivating them. The advantage of this approach is that it may be beneficial to improve the accuracy of genome binning (because different families of microorganisms were mixed in the co-culture) and to the subsequent preservation and genetic modification of the strains. However, this may also lead to the loss of some potentially non-cultivable microorganisms who may secrete antibiotics in biofilms.

## Materials and Methods

### Soil and strains

Soil samples were collected from two paddy fields that are located in Dajin Town, Wuxue City, Hubei province, China (115°33’E, 29°51’N). This area has a subtropical monsoon climate. In each paddy, five samples were taken from the top layer of soil, which is approximately 5-15 cm deep, and thoroughly mixed. After they were sampled, soils were partially air-dried and passed through a 2 mm sieve for use in subsequent experiments. The strains used in this study (Table S1) were all isolated from cultivated biofilm communities using a previously described method (23). The 16S rRNA genes of these strains were amplified with primers 27F and 1492R and sequenced using an ABI 3730XL DNA Analyzer (Applied Biosystems, US). The sequences were assembled by ContigExpress software (Invitrogen) and submitted to SINA Alignment Service to obtain their taxonomic classifications (39).

### Biofilm cultivation in a parallel flow system

The system was set up as previously described (23). For each sample, sieved soil (150 g, equivalent to 97.6 g dry weight) was mixed with 300 mL of sterilized ddH_2_O in a large beaker. The slurry was mixed at 150 rpm for 30 min, followed by gentle agitation at 30 rpm overnight. Microscope glass slides were covered with 1% autoclaved noble agar (Difco, BD, New Jersey, USA) several times and left until the agar cover solidified. The glass slides were then placed in a rack attached to a beaker containing soil slurry so that the slides were covered by the liquid phase of the slurry but did not disturb the solid. The beaker was incubated with shaking (30 rpm) for 7 days at room temperature. The glass slides were then removed from the beaker and briefly washed with 800 μL of phosphate buffer solutions and the agarose gels on the glass plates (i.e., biofilms) were collected into a tube. Then, 300 μL of sterilized glycerol was added to the tube, the biofilms were thoroughly mixed and stored at −80 °C to be used as a starter inoculum. The starter inoculum was inoculated onto the surface of seven agar-covered glass slides, which were then assembled between two holders in a homemade plexiglass flow chamber and left for 4 h to allow the bacteria to attach to the agar. After bacterial attachment, ~300 mL of autoclaved soil water (ASW) was circulated through silicone tubes over the slides in a closed circuit with a flow rate of 7.5 mL/min at room temperature for 7 days. Biofilms on glass slides were harvested at days 1, 2, 4 and 7 by removing one glass slide from the flow chamber. Slides with attached biofilms were rinsed twice with 0.9% NaCl, and biofilms were collected into a tube and divided equally into two parts. One part was subjected to 16S rRNA gene sequencing, and the other was used for strain isolation.

### DNA extraction and high-throughput 16S rRNA gene sequencing

Microbial DNA was extracted from the agarose surface that had been gathered from glass plates on different days using an E.Z.N.A.^®^ Soil DNA Kit (Omega, Norcross, GA, USA) according to the manufacturer’s instructions. The V4-V5 region of the bacterial 16S ribosomal RNA gene was amplified by PCR (95 °C for 2 min, followed by 25 cycles at 95 °C for 30 s, 55 °C for 30 s, and 72 °C for 30 s and a final extension at 72 °C for 5 min) using primers 515F 5’-barcode-GTGCCAGCMGCCGCGG)-3’ and 907R 5’-CCGTCAATTCMTTTRAGTTT-3’, where barcode is an eight-base sequence unique to each sample. PCR reactions were performed in triplicate in a 20 μL mixture containing 4 μl of 5 × FastPfu Buffer, 2 μL of 2.5 mM dNTPs, 0.8 μL of each primer (5 μM), 0.4 μl of FastPfu Polymerase, and 10 ng of template DNA. Amplicons were extracted from 2% agarose gels and purified using an AxyPrep DNA Gel Extraction Kit (Axygen Biosciences, Union City, CA, U.S.) according to the manufacturer’s instructions and were quantified using a QuantiFluor™ -ST (Promega, U.S.). Purified amplicons were pooled to be equimolar and paired-end sequenced (2 × 250) on an Illumina MiSeq platform according to standard protocols. Raw FASTQ files were demultiplexed and quality-filtered using QIIME (version 1.9.1) with the following criteria: (i) The 300 bp reads were truncated at any site receiving an average quality score <20 over a 50 bp sliding window, and any reads shorter than 50 bp were also truncated. (ii) Exact barcode matching was required, a two nucleotide mismatch was allowed in primer matching, and reads containing ambiguous characters were removed. (iii) Sequences that only overlapped more than 10 bp were assembled according to their overlap sequence. Reads that could not be assembled were discarded. Operational Taxonomic Units (OTUs) were clustered with 97% similarity cutoff using UPARSE (version 7.1 http://drive5.com/uparse/), and chimeric sequences were identified and removed using UCHIME. The taxonomy of each 16S rRNA gene sequence was analyzed by RDP Classifier (http://rdp.cme.msu.edu/) against the Silva (SSU123)16S rRNA database using a confidence threshold of 70% (40). Principal component analysis and temporal analysis were performed by STAMP software (version 2.1.3) (41). During temporal analysis of soil biofilms, an analysis of variance (ANOVA) test was employed, and the most significantly changed bacterial groups were identified with p-values < 0.05. The raw reads of 16S rRNA gene sequencing were deposited into SRA database (https://trace.ncbi.nlm.nih.gov/Traces/sra/sra.cgi?) under the BioProject of PRJNA420805.

### Crystal violet assay

A modified version of the crystal violet (CV) assay for the detection of biofilms using 96-well cell culture plates was applied as previously described (42). Biofilms formed on pegs of the Nunc-TSP lid system were measured after 24 h of incubation unless otherwise indicated. To wash off planktonic cells, the peg lid was successively transferred to three microtiter plates containing 200 μL of distilled deionized water per well. After the biofilms were stained for 20 min with 180 μL of a 0.1% (*w/v*) aqueous CV solution, the lid was again rinsed three times and then placed into a new microtiter plate with 200 μL of anhydrous ethanol in each well for 30 min to allow the stain to dissolve into the ethanol. The absorbance of CV at 590 nm was then determined by using a Multiskan™ GO Microplate Spectrophotometer (Thermo Scientific). Statistical analyses were conducted using an ANOVA test (OriginPro version 9.3, OriginLab Corp.) A p-value of less than 0.05 indicated a statistically significant difference.

### Meta-transcriptomic analysis

Exponential-phase cultures of *Novosphingobium sp*. G5, *Pseudomonas sp*. G7, *Flavobacterium sp*. G16 and *Microbacterium sp*. D12 were adjusted to an optical density of 0.15 at 600 nm in tryptic soy broth (TSB) medium. One milliliter of each culture and a mixture of containing 250 μL of each individual culture, were then inoculated into 50 mL of fresh TSB medium. After 12 h of incubation at 25 °C with shaking (200 rpm), cells were collected by centrifugation at 4000× g. Then, equal amounts of monoculture cells were mixed and treated as a monoculture control (CK). Total RNA of CK and the four-species co-culture were extracted with TRIzol reagent (Invitrogen) following the manufacturer’s instructions. RNA-seq libraries constructed with co-culture RNA and the control RNA were sequenced by HiSeq 2500 platform using standard protocols (BGI, China). After filtering, the remaining clean reads were *de novo* assembled into transcripts using Trinity (version 2.0.6) (43), followed by gene family clustering with Tgicl (version 2.0.6) (44) to get final unigenes. Binning of unigenes was performed by analyzing the taxonomic classification of BLAST hits in MEGAN (version 6.4.15) under the family level (45), as the four species in the co-culture are from four different families. MetaGeneMark (version 3.25) was employed to predict the putative genes in those unigenes (46). Reads were then separately mapped to the predicted genes of each species using bowtie2, differentially expressed genes were characterized by RSEM-EBSeq pipeline (version 1.2.22) (47, 48). This pipeline is capable of obtaining high confidence results in the absence of experimental replicates, and thus is very suitable for high-throughput screening applications. COG annotation of genes was carried out by aligning them to the COG database using BLAST (access Jan, 2017) (49). Raw sequence data were deposited in the SRA database (https://trace.ncbi.nlm.nih.gov/Traces/sra/sra.cgi?) under the BioProject of PRJNA401492.

### Whole genome sequencing of *Pseudomonas* sp. G7

The *Pseudomonas* sp. G7 strain genome was sequenced using a PacBio RS II platform and Illumina HiSeq 4000 platform at the Beijing Genomics Institute (BGI, Shenzhen, China). Four SMRT cells were used by the PacBio platform to generate the subreads set. PacBio subreads with length less than 1 kb were removed. The program Pbdagcon (https://github.com/PacificBiosciences/pbdagcon, accessed Feb, 2018) was used for selfcorrection. Draft genomic unitigs, which are uncontested groups of fragments, were assembled using the Celera Assembler against a highquality corrected circular consensus sequence subreads set. To improve the accuracy of the genome sequences, GATK (https://www.broadinstitute.org/gatk/, accessed Feb, 2018) and SOAP tool packages (SOAP2, SOAPsnp, SOAPindel, accessed Feb, 2018) were used to make single-base corrections. To trace the presence of any plasmid, the filtered Illumina reads were mapped using SOAP to the bacterial plasmid database (http://www.ebi.ac.uk/genomes/plasmid.html, accessed July 8, 2016). The complete genome sequence of *Psuedomonas* sp. G7 was submitted to NCBI Genbank database and annotated by Prokaryotic Genome Annoation Pipeline (accessed Mar 9, 2018) (50). To reveal the antibiotic biosynthesis gene clusters in the strain, the sequence was analyzed by antibiotics and secondary metabolite analysis shell (antiSMASH, version 4.0) (9), followed by manually correction.

### Comparative genomics

We used comparative genomics to analyze the sequence diversity of the two antibiotic synthesis BGCs found in *Pseudomonas* sp. G7. Briefly, the nucleotide sequences of two NPRS antibiotic BGCs were used to query the non-redundant nucleotide collection (nt database, accessed Apr 4, 2018) for any known similar BGCs by NCBI BLAST. As a result, five *Pseudomonas* spp. were found have an architecture similar to that of the two BGCs (Table 2). The genomes of the five species were then analyzed by antiSMASH (9), and the corresponding BGCs were identified. To compare BGCs, the protein sequences of their encoded genes were aligned and the phylogenic distance calculated by msa package (version 1.12.0) (51), then a phylogenetic tree was constructed using a maximum likelihood method with ape package (version 5.1) (52). Trees and distances were visualized using ggtree (version 1.12.0) (53) and pheatmap (version 1.0.8, https://CRAN.R-project.org/package=pheatmap), respectively.

### RNA extraction, cDNA synthesis and Quantitative real-time PCR (qRT-PCR)

The experiments were carried out as previously described (54). Briefly, RNA was isolated from 1 mL of broth cultures with TRIzol Reagent (Invitrogen) following the manufacturer’s instructions. For reverse-transcription PCR, the extracted RNA was first digested by gDNA wiper (included in the kit) to remove possible genomic DNA contamination and then used as a template for synthesis of cDNA using 6-mer random primers and the HiScript II 1st Strand cDNA Synthesis Kit (Vazyme). The cDNA was used to amplify the core synthetic genes of the NRPS BGC and the 16S rRNA gene of *Pseudomonas* sp. G7 using their specific primers with AceQ^®^ qPCR SYBR^®^ Green Master Mix (Vazyme) and the standard SYBR protocol in a Quantstudio 6 Flex instrument (Applied Biosystems). For each PCR, the experiments were performed in at least triplicate and the amplifications were checked by navigating the amplification curve and melting curve. Gene expression was normalized to the levels of 16S rRNA and the changes in gene expression were calculated using the 2^-ΔΔCt^ method (55).

## Data accessibility

16S rRNA gene sequences of soil biofilm isolates: Genbank accession MF689010-MF689033. 16S rRNA gene amplicon sequencing reads were submitted to SRA database under the BioProject of PRJNA420805. Meta-transcriptome sequencing reads were submitted to SRA database under the BioProject of PRJNA401492. Whole genome sequencing results were submitted to Genbank under the accession of CP027561.

## Statistics analysis

In addition to the above described statistical analysis with specific software, some other statistical analysis was performed with R (https://www.r-project.org/).

## Conflicts of interest

The authors declared there is no conflict of interest.

## Acknowledgements

This work was supported by the National Key R&D Program of China (2016YFD0800206), the National Natural Science Foundation of China (41522106). The authors would like to thank Dr. Qian Wang, currently works at Montana State University, for her inspirational comments on this study.

## Author Contributions

P.C. conceived the topic. C.H.G. and Z.L. performed experiments, C.H.G., Y.W., Q.H. and P.C. analyzed and interpreted the data. C.H.G. and P. C. wrote this paper with the help of all authors.

## Supplementary materials

Supplementary Figure S1: Alpha diversity of soil biofilm communities.

Supplementary Figure S2: The biofilm formation of mono-culture of seven soil isolates.

Strains: 1: *Aquitalea* sp. G2; 2: *Novosphingobium* sp. G5; 3: *Pseudomonas* sp. G7; 4: *Flavobacterium* sp. G16; 5: *Acinetobacter* sp. G18; 6: *Delftia* sp. D10; 7: *Microbacterium* sp. D12.

Supplementary Figure S3: Colony inhibitory assays. From left to right and from top to bottom, 3 microliters of bacterial culture from 7 different isolates were sequentially inoculated on LB plates (A) or on bacterial lawn of each of the 7 isolates (B-H) (as listed and ordered by 1-7 in Fig. 4). After 24 hours of cultivation, if the colonies can produce antibiotics and inhibit the growth of the lawn species, a inhibitory zone will be seen.

Supplementary Figure S4: Alignment of *Pseudomonas* sp. G7 Unigenes to *Pseudomonas* sp. DR 5-09 bananamide biosynthesis gene cluster.

Supplementary Table S1: Bacterial isolates from cultivated biofilm community.

Supplementary Table S2: Biofilms formed by single species and combinations (four strains).

